# Evolutionary Distance of Gene-Gene Interactions: Estimation under Statistical Uncertainty

**DOI:** 10.1101/2020.03.08.982710

**Authors:** Xun Gu

## Abstract

Consider the functional interaction of gene *A* to an interaction subject *X*; for instance, it is the gene-gene interaction if X represents for a gene, or gene-tissue interaction (expression status) if X for a tissue. In the simplest case, the status of this *A-X* interaction is *r*=1 if they are interacted, or *r*=0 otherwise. A fundamental problem in molecular evolution is, given two homologous (orthologous or paralogous) genes *A* and *B*, to what extent their functional overlapping could be by the means of interaction networks. Given a set of interaction subjects (*X*_1_, … *X*_*N*_), it is straightforward to calculate the interaction distance (*I_AB_*) between genes *A* and *B*, by a Markov-chain model. However, since the high throughput interaction data always involve a high level of noises, reliable inference of *r=*1 or *r*=0 for each gene remains a big challenge. Consequently, the estimated interaction distance (*I_AB_*) is highly sensitive to the cutoff of interaction inference which is subject to some arbitrary. In this paper we will address this issue by developing a statistical method for estimating *I_AB_* based on the *p*-values (significant levels). Computer simulations are carried out to evaluate the performance of different *p*-value transformations against the uncertainty of interaction networks.

## Introduction

Thanks to the growing amount and quality of omics data, the assembly of biological networks has played an important role to unveil the underlying cellular processes and evolution. In this scenario, Protein-Protein Interactions (PPIs) (Mosca et al. 2013; Hao et al. 2016) and Gene Regulation Networks (GRN) (Halton 2017) are among the most important and widely studied networks. A PPI networks is described in terms of proteins (nodes) and their physical/functional interactions (edges) (Nibbe et al. 2010; Sharan et al. 2007; Procaccini et al. 2016; Gustafsson et al. 2014), while a GRN network described in terms of gene-protein interactions. Note that transcriptome (RNA-seq) can be also viewed as the gene-tissue interactions. Both PPI and GRN networks are analyzed through the identification of subnetworks, or modules, showing specific topological and/or functional characteristics (Barabási et al. 2011; Vella et al. 2017; Hartwell et al 1999; Gursoy et al. 2008; Fraser 2005). For instance, a PPI module represents a group of proteins taking part in specific, separable functions such as protein complexes, metabolic pathways or signal transduction systems.

Here we focus on the evolution of interaction network. Initiated by an influential, but controversial study (Fraser et al. 2002) on the effect of protein-protein interactions on protein sequence evolution, interactivity at the DNA, protein and genetic levels has been the major topic in the study of systems biology and evolution (von Mering et al. 2002; Wagner 2001; 2003; Jordan et al. 2003; Kim WK, Marcotte 2008; Sun and Kim 2011). Given the network sizes, typically involving thousands of elements, it often requires in-silico automated methods (Ma’ayan 2008; Grindrod, P. & Kibble 2004). However, it has been shown that the interaction inferences based on large-scale omics data are subject to a high level of statistical noises (Benjamini and Hochberg 1995; Shaffer 1995; Benjamini and Yekutieli 2001; Efron et al. 2001; Efron 2004; Smyth 2004). Consider the functional interaction of gene *A* to any other gene *X*. In the simplest case, the status of this *A-X* interaction is *r*=1 if these two genes are connected, or *r*=0 otherwise. For two duplicate genes *A* and *B*, an important measure for their functional overlapping is the number (*n_11_*) of other (*X*) genes that interact with both duplicate genes *A* and *B* (*A-X* and *B-X*). One may utilize a simple Poisson model to estimate the interaction distance (*D_I_*) between duplicates, i.e., the average number of interaction changes (losses and gains). For a set (*n*) of genes, each of which interacts with *A*, or *B* or both, one can calculate the proportion of interaction divergence *q*=1−*n*_11_/*n*, and estimate the interaction distance by *D_I_*=−*n* ln (1−*q*). However, since the high throughput functional genomic data involve high level noises, statistical evaluation of inferred interactions becomes a big challenge in the genomic analysis. We shall address this issue, i.e. how to take the statistical uncertainty in the evolutionary analysis.

## New Methods

### Statistical representation for gene-gene interactions

#### p-Value presentation for single gene-query interaction

Many genomic studies have adopted a *p*-value to characterize the statistical significance of any gene-query interaction. That is, instead of a binary (*r*=1 or 0) status of any gene-query (*A-X*) interaction, a *p*-value is assigned for the interaction between a gene and a query; a small *p*-value, e.g., *p*=0.001, means that the interaction is highly statistically significant, and *vice versa*.

#### Multiple-test problem

Usually *p*-values of high throughput gene-gene interactions are subject to the sophisticated multiple-test problem (Benjamini and Hochberg 1995; Benjamini and Yekutieli 2001). Intuitively speaking, it depends on a pre-specified cut-off for the *p*-value, that is, the interaction status is assigned as ‘1’ if the *p*-value is less than the cutoff; otherwise the interaction is null. How to determine the cut-off is controlled by the false positive rate of discovery, usually called the *q*-value for a given set of interactions. A *q*-value, say 0.1, means that for all gene-gene interactions predicted as ‘active’, the proportion of false-positive cases has been set to be 10%. To calculate the *q*-value under a given p-value cut-off, we have to model the *p*-value distributions under the null and the alternative hypotheses, respectively (Efron 2004; Erron et al. 2001).

#### Challenge for evolutionary analysis of interactions

While it is widely accepted that the evolutionary distance of interactions between homologous genes is important for understanding the pattern of genome evolution, the estimation problem has not been well addressed. Suppose that we have two genes (*A* and *B*) that have *p*-values to *N* interaction queries (*X*_*1*_,‥, *X*_*N*_), respectively. Apparently, the distance estimation becomes straightforward if all states of *A*-*X_i_* and *B-X_i_* (*i*=1,…, *N*) are unambiguously observed. As discussed above, statistical inference of interaction status for a set of *p*-values is controlled by the false-positive rate (the *q*-value). As a result, the estimated interaction distance is highly sensitive to the *q*-value cutoff selected.

### A general *p*-value framework for the interaction evolution

Our goal is to develop a practically feasible approach to overcome this problem by treating these *p*-values as observations. We use the statistical approach for modeling *p*-values similar to that used in the analysis of multi-test problem. Instead of determining the false positive rate for a given set of predictions, we combine the *p*-value model and the underlying evolutionary model to develop a cutoff-free method for estimating the interaction distance.

#### Expression distance defined by transformed p-values

For *p*-value presentation of gene-query interactions, we have to develop an explicit model for the interaction evolution (Fig.1). Consider two homologous genes *A* and *B* with an interaction query *X*, there are four combined patterns (*r*_*A*_, *r*_*B*_): (1, 1), (1, 0), (0, 1) and (0, 0), respectively. For instance, (1, 1) means that both genes have the interaction with the same query *X*; (1, 0) means gene *A* has the interaction but gene *B* does not, and so forth. The probability of each pattern is denoted by *P*(*r*_*A*_, *r*_*B*_), where *r_A_, r_B_*=1 or 0, respectively.

**Fig. 1.**
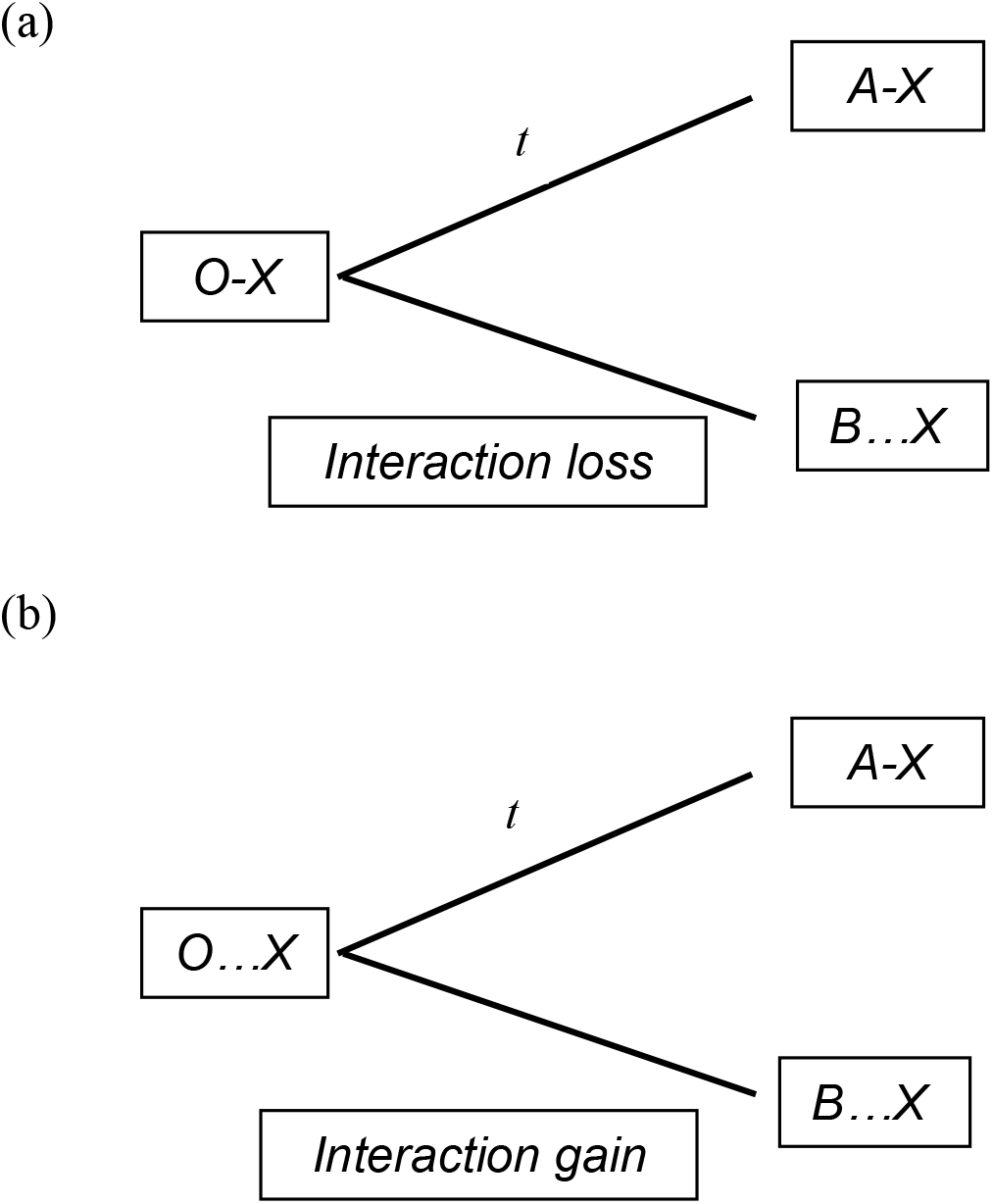
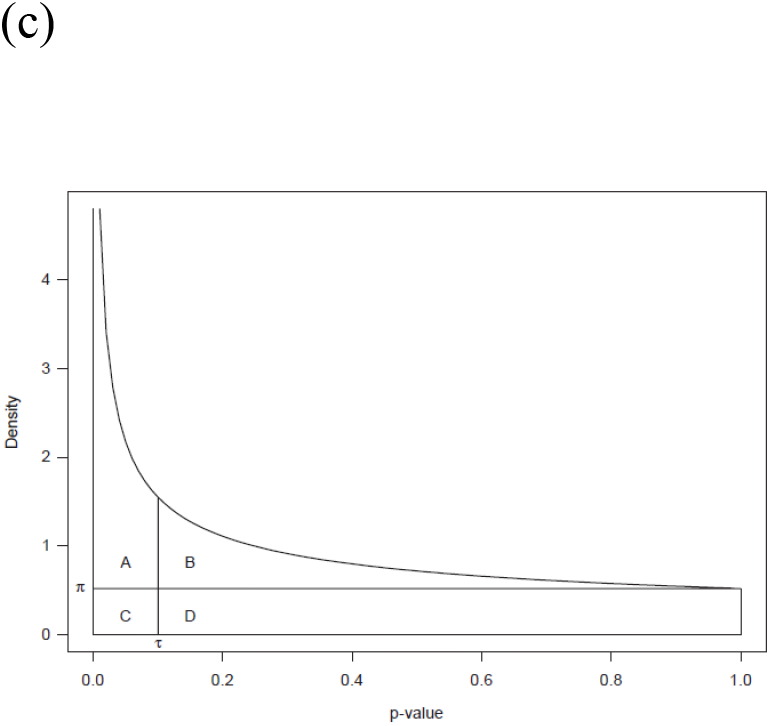
Schematic illustration of interaction evolution. Suppose two homologous genes (*A* and *B*) diverged *t* time units ago, via either speciation or gene duplication. For a given interaction query gene (X), gene A has an active interaction with X (solid line) whereas gene B has no interaction with X (dashed line). There are two possibilities for their common ancestor (gene O): in the case of active interaction between O and X, there is an interaction loss in the B-lineage (panel a), otherwise there is an interaction gain in the A-lineage (panel b). (c) Schematic illustration of the BUM distribution for *p*-values. Region A corresponds to the occurrence of true positives; region B corresponds to the occurrence of false negatives; Region C corresponds to the occurrence of false positives and region D corresponds to the occurrence of true negatives.

Let *p*_*A*_ and *p*_*B*_ be the *p*-values for interactions *A-X* and *B-X*, respectively. Let *y_A_* and *y_B_* be any given transformations of *p*_*A*_ and *p*_*B*_, respectively. Two simple forms that are practically useful are *y*=−ln (*p+p_0_*) and *y=p*, respectively (*p_0_* is the small number to avoid *y* tends to be infinite when *p* approaches 0, usually one may choose *p_0_*=1/*M*; where *M* is the total number of interactions under study). Next, we define the expectation of squared *y*-score differences between genes *A* and *B* as follows

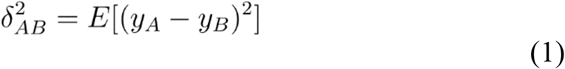

where *E* is for taking expectation.

While it is straightforward to estimate δ^2^_*AB*_ from the interaction data as long as the transformation formula is specified, the challenge is how to connect δ^2^_*AB*_ to the evolutionary model of interactions. Our idea is to expand the right hand of Eq.(1) by the means of conditional expectations with respect to interaction patterns (*r*_*A*_, *r*_*B*_), which can be concisely written by

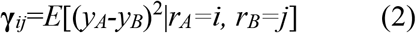

where *i, j*=0 or 1. Let *P*(*r*_*A*_, *r*_*B*_) be the probability of an interaction pattern (*r*_*A*_, *r*_*B*_). According to the probability theory, we have

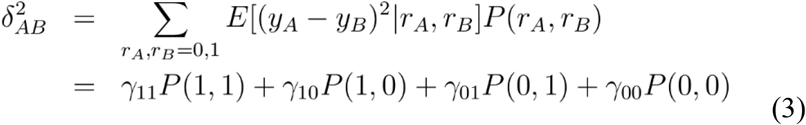

In short, according to Eq.(3), δ^*2*^_*AB*_ can be decomposed into two components: the first component is the interaction uncertainty measured by γ_*ij*_, and the second component is the evolution of interactions (gain or loss) measured by the probabilities *P*(*r*_*A*_, *r*_*B*_).

### Evolutionary model to determine *P*(*r*_*A*_, *r*_*B*_)

Suppose that gain and loss of interactions are the major mechanisms for the interaction evolution of a gene, which are independent of other genes. We then develop a simple Markov-chain model as follows. Let π_1_ be the probability of active interaction (*r*=1), and π_0_=1−π_1_ be that of inactive interaction (*r*=0). Let λ be the evolutionary rate of interaction. Under the Markov-chain model, the gain rate and the loss rate of an interaction are given by π_1_λ and π_0_λ, respectively. It follows that the transition probabilities from state *i* to state *j* after the *t* time units, *P_ij_*(*t*), *i, j*=0, or 1, are given by *P_11_*(*t*)= π_1_+π_0_*e^−λt^*, and *P_10_*(*t*)= π_0_(1−*e^−λt^*); similarly, we have *P_00_*(*t*)=π_0_+π_1_*e^−λt^*, and *P_01_*(*t*)= π_1_(1−*e^−λt^*). In the case of two homologous genes, the probability for any observed pattern *P*(*r*_*A*_, *r*_*B*_) can be calculated as follows

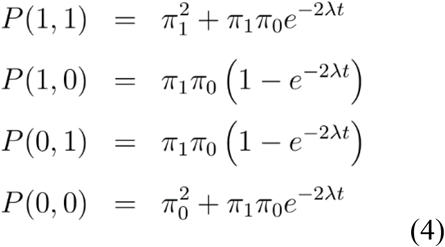

### A general formula for the evolutionary distance of interactions

When the probability of an interaction pattern (*r*_*A*_, *r*_*B*_), *P*(*r*_*A*_, *r*_*B*_), is determined by Eq.(4), we can derive a general formula for the evolutionary distance of interactions by combining Eq.(3) by Eq.(4), resulting in

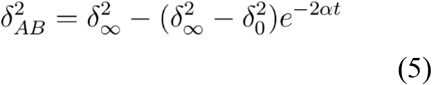

where δ^*2*^*∞* and δ^2^_*0*_ are given by

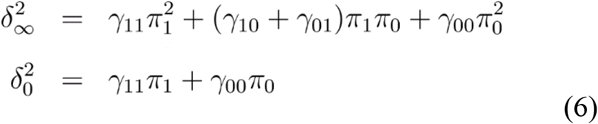

Eq.(5) shows that, when *t*=0, δ^2^*_AB_*=δ^2^_*0*_, and δ^2^_*AB*_ increases with *t* and ultimately reaches δ^*2*^_∞_ as *t*→∞. One may further define the effective proportion of different interactions between genes *A* and *B*

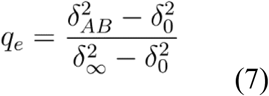

such that *q_e_* satisfies

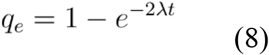

Then, given the number (*N*) of interaction queries, the interaction distance defined by *I_AB_*=2π_1_π_0_λ*t*, is given by

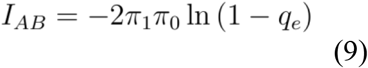

## Results and Discussion

Based on the theoretical framework formulated above, we develop a computational procedure that is suitable to a variety of OMICS data types.

### The beta-uniform mixture (BUM) distribution of *p*-values

While *p*-values arising from the null hypothesis are distributed uniformly on the interval (0, 1), those arising from the alternative hypothesis follows a distribution denoted by φ(*p*), which is generally modeled as a beta distribution. Therefore, the distribution of the set of *p*-values can be written as a beta-uniform mixture (BUM) consisting of a uniform (0, 1) component for the null hypothesis and the beta component for the alternative hypothesis, with the pdf given by

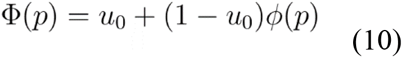

where *π_0_* is the proportion of null hypothesis (*r*=0); and the second term for the Beta component. Here we implement a special form of beta distribution by setting *β_0_*=1, resulting in

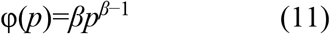

for 0*<p*≤1, 0*<π_0_<*1, and 0<*β*<1. As shown in Fig.1(c), Eq.(11) provides a reasonable model for the distribution of *p-*values arising from high throughput genomics (Storey and Tibshirani 2003). It is a curve that asymptotes at *x*=0 and monotonically decreases to its minimum of *π_0_*+(1−*π_0_)β* at *x*=1. This curve approximates the anticipated distribution of the *p*-values arising from a genomic experiment. Under the null hypothesis, the *p*-values will have a uniform density corresponding to a flat horizontal line. Under the alternative hypothesis, the *p*-values will have a distribution that has high density for small *p*-values and the density will decrease as the *p*-values increase. The overall distribution will be a mixture of *p*-values arising from the two hypotheses.

### Parameters of interaction uncertainty under the BUM model

Suppose that *y_A_* and *y_B_* are independent, which follow the same distribution conditional of *r*=0 or 1. Then **γ**_*rA, rB*_ defined by Eq.(2) can be further simplified as follows

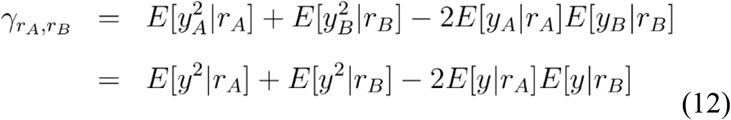

To calculate **γ**_*rA, rB*_, we have to consider the null hypothesis of *r*=0 (no interaction) and the alternative separately. Under the null hypothesis, the *p*-value follows a uniform distribution. For a given *p*-value transformation, *y*=*f*(*p*), let 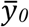 and *σ*_0_^2^ be the mean and variance of *y* under the null *r*=0, respectively, which are given by

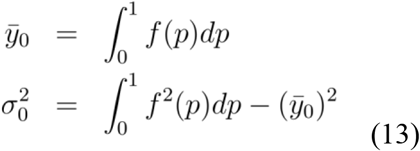

One may easily show that 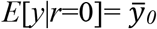 and 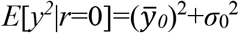. Under the alternative hypothesis of *r*=1, the distribution of *p*, denoted by φ(*p*), is usually modeled by a beta distribution given by Eq.(11). Similar to above, let 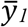 and *σ*_1_^2^ be the mean and variance of *y* under the alternative of *r*=1, respectively, as given by

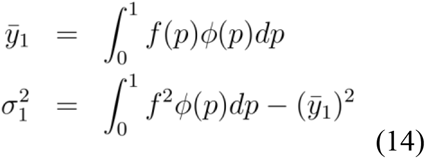

Obviously we have 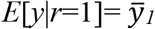 and 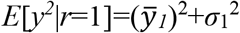. Putting together, those coefficients of interaction uncertainty (γ_*11*_, γ_*10*_, γ_*01*_ and γ_*00*_) defined by Eq.(12) are given by

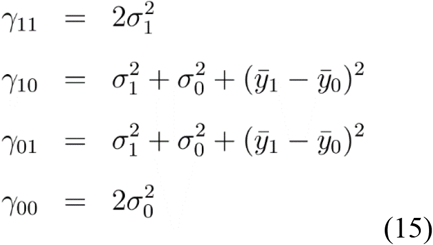

respectively.

### Types of *p*-value transformation

We shall implement a statistical procedure to estimate those parameters on the right hands of Eq.(15), as long as the *p*-value transformation is specified. In this study we consider two *p*-value transformations: the *p*-based, and the −ln(*p*+1/*M*)-based, where *M* is the total number of interactions.

#### The p-based method

The simplest one is to use the *p*-value directly, i.e., *y=p*. Note that under the null of no interaction (*r*=0), *p* follows a uniform distribution in [0,1] with the mean 1/2 and the variance 1/12. In this case, Eq.(15) can be simplified as follows

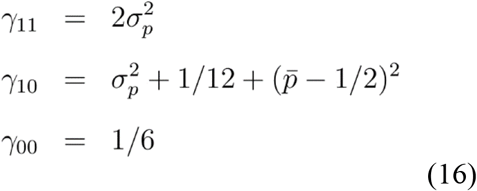

where 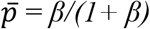 and 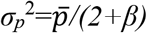 are the mean and variance of *p*-values under the beta distribution of Eq.(11).

#### The −ln(p+1/M)-based method

In spite of simplicity, we have realized that *y=p* transformation may be subject to some sampling problems. For instance, an interaction associated with a *p*-value of 0.001 is statistically sound, while that with more than 0.05 is usually considered as non-significance. Consider a hypothetical example that for two duplicate genes *A* and *B*. Assume *p*_*A*_=0.001 and *p*_*B*_=0.5 for the functional interactions A-X and B-X, respectively, so that (*p_A_−p_B_*)^2^=0.4999^2^. Secondly, for another interaction pair A-X′ and B-X′, we assume that the *p*-values are *p′_A_*=0.30 and *p′_B_*=0.8 so that (*p′_A_-p′_B_*)^2^=0.5^2^. The virtually same score between two cases is apparently counter-intuitive, because one may statistically infer that gene *A* interacts with *X* but not for gene *B*, whereas both *A* and *B* are unlikely to interact with gene *X′*.

We try to use a (negative) log-transformation score (*y*) for the *p*-value of an interaction, i.e., *y=−ln p*, to avoid this difficulty. In the above case, we observed that (*y_A_−y_B_*)^2^=6.22^2^ (for A-X and B-X) is much higher than (*y′_A_−y′_B_*)^2^=0.98^2^ (A-X′ and B-X′), which makes much more sense. While this log-transformation can effectively reduce the random effects, one problem is that *y=−ln p* does not converge when *p* approaches to zero, which may cause a considerable (upward) sampling bias. We thus recommend a simple correction by adding a pseudo-count, that is, *y*=−ln(*p*+1/*M*), where *M* is the total number interactions under study.

Since *M* is usually large, one can show that under the null hypothesis *y* approximately follows an exponential distribution with the mean 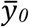 and variance *σ_0_*^2^. Together, we have

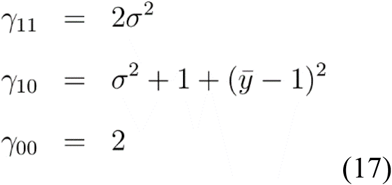

where 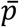 and *σ*^2^ are the mean and variation of *y*-values under the alternative hypothesis of *r*=1, respectively.

### Computational procedure and implementation

Under the *R*-environment, we implement the following computational procedure to estimate the evolutionary distance of interactions based on *p*-values.

i. Given high throughput *p*-values, we use several statistical methods to estimate *π_0_*, the proportion of null hypotheses of interactions. First, the LOESS-based method (Pounds and Cheng 2004) applies LOESS to the *p*-value spacing to obtain an estimate of PDF, and takes the minimum value of the estimated PDF as an estimate of *π_0_*. Second, the CDF-based method (Cheng et al. 2004) uses an estimator for the CDF (Cumulative Distribution Function) of p-values in the form of a B-spline series with strategically designed knot sequence to achieve a desirable shape for the *p*-value cumulative distribution. The PDF is simply the first derivative of the CDF and the minimum of PDF is taken as an estimate of *π_0_*. Third, exploiting the fact that p-values from true null hypotheses are uniformly distributed, the asymptotic uniform method (Storey and Tibshirani method 2003) uses a tunable estimate, that is, *π_0_*(*c*)= #{*p*_i_>*c*;i=1,…*M*}/*M*(1-*c*), with *c* as the tuning parameter. It then fits a natural cubic spline with 3 degrees of freedom to the data of *π_0_*(*c*) on a series of values of *c* such as *c*=0, 0.05,…, 0.90, and finally the value of the fitted spline line at the end point *c*=1 is taken as the estimate of *π_0_*. The technical details of these methods can be found in the original publications.
ii. Treating the estimate of *π_0_* as known, the quasi-maximum likelihood estimate of parameter *β* can be obtained by fitting the beta-uniform mixture (BUM) model in Eq.(11) to the observed *p*-values.
iii. Calculation of γ_*11*_, γ_*10*_, γ_*01*_ and γ_*00*_ numerically for *y=p* and *y*=ln(*p*+1/*M*), respectively, leading to the estimation of δ^*2*^_∞_ and δ^*2*^_*0*_ by Eq.(6).
iv. Given the sample size (*N*) of interaction queries, δ^*2*^_*AB*_ can be calculated as follows

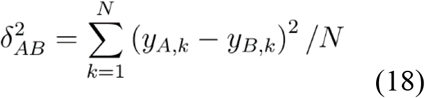

The effective proportion of different interactions between genes (*q_e_*) can be estimated according to Eq.(7). Then the evolutionary distance of interactions distance can be estimated by Eq.(9). In short, estimation of *I_AB_* turns out to estimate the effective proportion of different interactions between genes, which can be achieved when the transformed *p*-value (the *y*-score) is specified.

### Simulation study

#### Statistical uncertainty of interactions

Experimental noises in high throughput genomics data could make any biological analysis unreliable. Hence we wish to investigate this effect on the estimation of interaction distance. Since the value of parameter *β* is inversely related to the level of experimental noise, we design the following simulation study pipeline. We choose *β=* =0.01, 0.05, 0.1, 0.3, 0.5, and 0.8. Intuitively, a low *β* value indicates a high number of biological replicates, and *vice versa*. According to Eq.(11), these values correspond to type-II error at the 0.05 significance level of 0.03, 0.14, 0.26, 0.59, 0.78, and 0.85, respectively, as calculated by 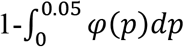.

#### Simulation theme of interaction evolution

Under the BUM model of interaction uncertainty, the simulation theme of interaction evolution is designed as follows. (*i*) Given the evolutionary scenario of two species with *N* interaction queries, simulate the interaction pattern according to the Markov-chain model. We set the interaction distance *I_AB_*=0.1, 0.3, 0.5, 0.8, or 1.0, respectively; the number of interaction queries is *N*=100, 200 or 500, respectively; and the proportion of active interaction is set to be *π_1_*=0.1, 0.3, 0.5, 0.7 or 0.8, respectively. (*ii*) In each case, we estimate those interaction uncertainty parameters for two *p*-value transformations, *y=p*, and *y*=−ln(*p*+1/*M*), respectively, as well as the evolutionary distance. And (*iii*) carry out 1000 simulation replicates and analyze the statistical properties in each case. The goal of our simulation study is to examine the systematic bias and sampling variance of *I_AB_* under various conditions (Table 1).

**Table 1.** A summary of simulation studies to evaluate the statistical performance of the interaction distance.

|  | *Distance ($I_{AB}$)* |  |  |
| --- | --- | --- | --- |
| $y$ | $p$ | $-\ln p$ | $-\ln (p+1/M)$ |
| $\beta=0.8$ | | | |
| $I_{true}= 0.1$ | 0.033±0.503 | 0.105±0.043 | 0.106±0.024 |
| $I_{true}= 0.5$ | 0.409±0.204 | 0.477±0.050 | 0.475±0.053 |
| $I_{true}= 1.0$ | 0.845±0.308 | 0.945±0.102 | 0.986±0.103 |
| $\beta=0.5$ | | | |
| $I_{true}= 0.1$ | 0.087±0.107 | 0.107±0.023 | 0.103±0.022 |
| $I_{true}= 0.5$ | 0.468±0.103 | 0.568±0.043 | 0.511±0.040 |
| $I_{true}= 1.0$ | 0.944±0.308 | 1.544±0.603 | 0.974±0.407 |
| $\beta=0.10$ | | | |
| $I_{true}= 0.1$ | 0.092±0.036 | 0.122±0.036 | 0.102±0.017 |
| $I_{true}= 0.5$ | 0.485±0.132 | 0.582±0.315 | 0.512±0.036 |
| $I_{true}= 1.0$ | 0.968±0.167 | 1.868±0.804 | 0.968±0.042 |

Note: In each case, the number of query genes ( $N$ ) is set to be 500 and $\pi_I=\pi_O=0.5$ .
Sampling variances are calculated based on 1000 simulation replicates.

#### Effects of p-value transformation

When the simplest *y=p* transformation is used, our simulation shows a large sampling variance and severe underestimation bias, especially in those cases when the parameter *β* is set be large (>0.5). Indeed, a small *β* value (<0.05) can effectively reduce both sampling variance and bias. Note that a low *β* value indicates a high number of biological replicates, our observation suggests that the performance of *y=p* transformation for the estimation of *I_AB_* is reasonable acceptable only when the number of biological replicates is sufficient. Moreover, for closely related species when the number of alternative hypotheses is small, estimation of *I_AB_* becomes biologically meaningless when the network uncertainty is overwhelming (a large *β* value) (Table 1). These results, together, suggest that, using the p-value directly without any transformation is not recommended, because the estimated evolutionary distance of interactions could be highly sensitive to the network uncertainty.

By contrast, the −ln(*p+1/M*)-distance shows some nice statistical properties in the case of low biological replicates; both sampling variance and estimation bias are generally acceptable. We have examined the effect of the pseudo-count (1/*M*). Indeed, the −ln(*p*)-distance becomes statistically unreliable in the case of small *β* value (<0.05), especially when the number of query genes *N* is small. This observation can be explained as some genes may receive very low *p*-values (very high significance levels), resulting in some extremely high *y*=−ln(*p*) values. Fortunately, this defect can be effectively corrected by the −ln(*p+1/M*) distance. While the −ln(*p+1/M*) distance overall performs satisfactory, it tends to underestimate the distance when the number of query genes is small, such as less than 200.

#### Effects of evolutionary parameters

First, the interaction distance estimation *I_AB_*, as expected, is asymptotically unbiased, i.e., the estimate *I_AB_* is statistically unbiased for sufficiently large sampling size of interaction queries (*N*). Similarly, the sampling variance of *I_AB_* decreases with the sampling size of interaction queries (*N*). Second, the asymptotic rate is dramatically affected by the parameter *β* in the BUM model. It becomes very slow when *β* is large, suggesting that an accurate estimation of interaction distance becomes difficult when the experimental noise is high. Third, the proportion of NA (not applicable) cases increases with the increase of *β*, which makes the distance estimation practically not useful particular when the sampling size (*N*) is small. Forth, as a general tendency, the distance estimation usually has nice statistical properties in the cases around *π_1_*=*π_0_*=0.5.

### Outlook for further study

Our further study will be focused on how to improve the efficiency and applicability of the proposed method. Several research lines are considered. For instance, we try to find the *p*-value transformation such that it not only can optimize statistical properties but also biologically interpretable. In the case of multiple genes that may represent a gene family evolution or species evolution, one may develop a distance-based approach to investigating the evolutionary pattern of interactions. Yet, it remains a challenge to develop a generalized likelihood function of interactions under a phylogeny when the interaction uncertainty is taken into consideration.

